# Estimating Microbial Interaction Network:Zero-inflated Latent Ising Model Based Approach

**DOI:** 10.1101/2020.06.02.130914

**Authors:** Jie Zhou, Weston D. Viles, Boran Lu, Zhigang Li, Juliette C. Madan, Margaret R. Karagas, Jiang Gui, Anne G. Hoen

## Abstract

**Motivation:** Throughout their lifespans, humans continually interact with the microbial world, including those organisms which live in and on the human body. Research in this domain has revealed the extensive links between the human-associated microbiota and health. In particular, the microbiota of the human gut plays essential roles in digestion, nutrient metabolism, immune maturation and homeostasis, neurological signaling, and endocrine regulation. Microbial interaction networks are frequently estimated from data and are an indispensable tool for representing and understanding the relationships among the microbes of a microbiota. In this high-dimensional setting, the zero-inflated and compositional data structure (subject to unit-sum constraint) pose challenges to the accurate estimation of microbial interaction networks.

**Method:** We propose the *zero-inflated latent Ising* (ZILI) model for microbial interaction network which assumes that the distribution of relative abundance of microbiota is determined by finite latent states. This assumption is partly supported by the existing findings in literature [20]. The ZILI model can circumvents the unit-sum constraint and alleviates the zero-inflation problem under given assumptions. As for the model selection of ZILI, a two-step algorithm is proposed. ZILI and two-step algorithm are evaluated through simulated data and subsequently applied in our investigation of an infant gut microbiome dataset from New Hampshire Birth Cohort Study. The results are compared with results from traditional Gaussian graphical model (GGM) and dichotomous Ising model (DIS).

**Results:** Through the simulation studies, provided that the ZILI model is the true generative model for the data, it is shown that the two-step algorithm can estimate the graphical structure effectively and is robust to a range of alternative settings of the related factors. Both GGM and DIS can not achieve a satisfying performance in these settings. For the infant gut microbiome dataset, we use both ZILI and GGM to estimate microbial interaction network. The final estimated networks turn out to share a statistically significant overlap in which the ZILI and two-step algorithm tend to select the sparser network than those modeled by GGM. From the shared subnetwork, a hub taxon Lachnospiraceae is identified whose involvement in human disease development has been discovered recently in literature.

**Availability:** The data and programs involved in Section 4 and 5 are available on request from the correspondence author.

**Supplementary information:** Supplementary materials are available at *Bioinformatics*

## 1. Introduction

The human microbiome, the collection of trillions of microbial organisms that live in our body spaces, belong to one of thousands of different species [15, 24]. The organisms that inhabit the human gut are an additional source of genetic diversity that can influence metabolism and modulate drug interactions [45]. Recent advances in genomic technologies enable production of thousands of 16S rRNA sequences per sample [47] and are powerful tools to explore the basic biology about human microbiome. Nevertheless, analyzing microbiome data and converting them into meaningful biological insights are still challenging tasks. First, the observed absolute abundance in sequencing experiment cannot inform the real absolute abundance of molecules in the sample which can be attributed to the sequence depth associated with the sequencing experiment. Multiple normalization methods have been proposed in literature to solve this problem in which total sum scaling (TSS) is one of such methods that has been widely used in practice[2, 7, 22, 27, 37]. TSS scales each sample by the total read count and yields the relative abundance. However, the statistical analysis based on relative abundance can easily lead to spurious association due to the unit-sum constraint [1, 26, 28, 34, 44]. Further complicating the analysis of microbiome data is the zero-inflated distribution of read count [45]. As for the dataset in Section 5, among the 134 taxa, there are only 6 taxa whose nonzero observation proportions are greater than 80%. Zero inflation stems from the fact that the majority of the amplicon sequence variants (ASVs) either physically do not exist in the subject or are below the detection threshold for the given sample [24]. Another hurdle for analyzing the microbiome data is its high-dimensionality which usually involves hundreds of microbes and consequently models equipped for this modeling task should be employed.

Microbial interaction network (MIN) is an indispensable tool for representing and understanding the relationships among the microbes [12, 13, 15, 17]. Traditionally, the interactions are discovered through co-culture experiments which routinely involve only small number of species in an artificial community [21, 23]. Modern researches try to use the data from real environments such as human gut to infer the association among the microbes [3, 4, 5, 33]. Consequently, the corresponding statistical inferences of MIN have received much attention in recent years. However, the hurdles mentioned above hinder the effective estimation of MIN. As a compromise, most of the existing studies estimate the MIN under the oversimplified conditions [8, 29]. For example, the studies in [8] ignored the unit-sum constraint and only considered the microbes whose nonzero observation proportion being higher than a given threshold. In [29], in order to deal with the zero inflation, the authors pooled all the sparse taxa together and formed a composite taxon which was not sparse anymore.

In light of the difficulties in MIN estimation, in this paper we propose the zero-inflated latent Ising model (ZILI) to characterize the underlying data generation mechanism of microbiota. Latent models such as hidden Markov models, state space models *et al*. have been widely used in economics, engineering and biology *et al*. Despite their popularity across disciplines, latent models have not been investigated for microbiome data yet. Incidentally, the studies in [20] found that the microbiota in human vagina could be characterized by finite states which provided a simple and intuitive understanding about the MIN in vagina. Inspired by the work in [20], we assume in ZILI that each of the *p* microbes in a microbiota can be characterized by a latent random variable *Z_j_* (1 ≤ *j* ≤ *p*). While for the random vector **Z** = (*Z*_1_, …, *Z_p_*), the multiclass Ising model is employed to characterize the joint distribution of **Z**. The relative abundances for each microbe are assumed to come from a zero-inflated mixture distribution which depends on the realization of **Z**. Given such a modeling framework, we propose a two-step algorithm for the model selection of ZILI. Specifically, in first step we estimate the states for each component of **Z** by transforming the relative abundances into categorical data. This step is implemented by an efficient dynamic programming algorithm. Based on the estimated state, in second step we use *L*_1_-penalized group logistic regression to select the nonzero parameters involved in ZILI. Through simulated data, we investigate the performance of two-step algorithm and demonstrate its effectiveness when ZILI is the underlying data generation model. We employ the Gaussian graphical model (GGM) and dichotomous Ising models (DIS) to analyze the same simulated data which show little power to select the true model. We apply both ZILI and GGM to an infant gut microbiome dataset from the New Hampshire Birth Cohort Study. It turns out the networks estimated by ZILI and GGM share a statistically significant subnetwork and ZILI shows the tendency to select sparser network than GGM. Within the shared subnetwork, Lachnospiraceae is identified as the hub taxon. Recent researches have found that Lachnospiraceae widely exists in human gut [38] and is related to some severe diseases such as non-alcoholic fatty liver disease and inflammatory bowel diseases *et al* [35, 39]. Since this important taxon is identified by both models, this may indicate that both ZILI and GGM can explain part of the information encoded in the relative abundance and the ZILI model can serve as a competitive tool for the MIN selection.

The organization of this paper is as follows. In Section 2, the ZILI model is detailed. The related estimation procedures for ZILI are described in Section 3. Simulation studies are carried out in Section 4. Section 5 is devoted to compare ZILI and GGM through an infant gut microbiome dataset. Section 6 concludes with a review about ZILI model.

## 2. Zero-inflated Latent Ising Model for MIN

In this section, we introduce the zero-inflated latent Ising (ZILI) model for the microbial interaction network which provides an alternative way to handle the problem of unit-sum constraint and zero inflation. Suppose that there are *p* taxa in the microbiota of interest. For jth taxon (*j* = 1, …, *p*), let *Z_j_* denote its latent state variable which has the following multinomial distribution,

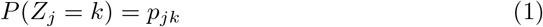

for *k* = 0,1, …, *K_j_* – 1 with 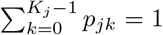. For example, there may be three states for *Z_j_* corresponding to three different states of relative abundance, (high, medium, low). This assumption can be partly justified by the existing findings in literature [20]. The studies in [20] found that the composition of vaginal bacterial communities can be characterized by five states. The microbiota for a given subject can be classified into one of these five states. The state may be affected by the exogenous factors such as sexual activity, menstruation *et al*. In order to study the general relationship among the microbiota, equation (1) generalizes the results in [20] and assumes there are finite states for each microbe. For ease of exposition, in the following we assume that all *Z_j_*’s are *K*-level variables. The arguments can be generalized to the more general situation straightforwardly for which *K_j_* may differ for different microbes. We pool all the *Z_j_*’s together and form the vector **Z** = (*Z*_1_, …, *Z_p_*) for which multiclass Ising model is employed to characterize its joint distribution,

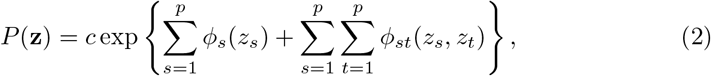

where *ϕ_s_* and *ϕ_st_* are the potential functions associated with *Z_s_* and (*Z_s_,Z_t_*) respectively. Since our aim is to estimate the conditional relationship among *Z_j_*’s, these potential functions can be parameterized as follows. For each 1 ≤ *s* ≤ *p*, and *l* ∈ {0, …, *K* – 1}, define *I*[*z_s_* = *l*] = 1 if *z_s_* = *l* and 0 otherwise. Then we have

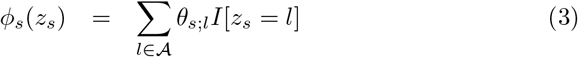

for *s* ∈ {1, 2, …, *p*} and 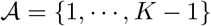 while

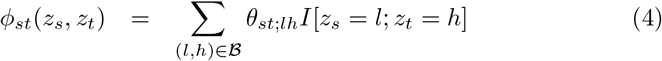

for (*s,t*) ∈ {1, …, *p*}^2^ and 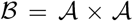. The unknown parameters in (3)-(4) include ***θ*** = {*θ*_*j*;*l*_, *θ*_*jt*;*lh*_ *j* = 1, …, *p*, *t* = 1, …, *p*, *j* ≠ *t*, *l* = 1, …, *K* – 1, *h* = 1, …, *K* – 1}.

Based on (2)-(4), for 1 ≤ *i* ≤ *n*, 1 ≤ *j* ≤ *p*, we have the following equation hold [36, 46],

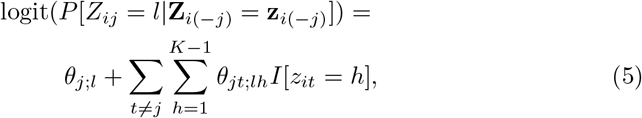

where **Z**_*i*(–*j*)_ = (*Z*_*i*1_, …, *Z*_*i*(*j*–1)_, *Z*_*i*(*j*+1)_, …, *Z_ip_*)^*T*^ with *Z_ij_* the *i*th observation of *Z_j_*. From (5), it can be shown that *θ*_*j*;*l*_ is the log-odds for event *Z_j_* = *l* given that the other *Z_t_*’s, *t* ≠ *j* are all zero. Similarly, *θ*_*jt*;*lh*_ is the log-odds ratio describing the association between events *Z_j_* = *l* and *Z_t_* = *h* given that all the other components of **Z** are fixed to zero. For more details about the interpretation of these quantities, see [36] and references there. Let ***θ***_*jt*_ = (*θ*_*jt*;11_, …, *θ*_*jt*;1(*K*–1)_, *θ*_*jt*;(*K*–1)1_, …, *θ*_*jt*;(*K*–1)(*K*–1)_)^*T*^. Vector ***θ***_*jt*_ reflects the relationship between *Z_j_* and *Z_t_*. If all the components of ***θ***_*jt*_ are zero, *Z_j_* and *Z_t_* turn out to be independent. If there exist nonzero components in ***θ***_*jt*_, then *Z_j_* and *Z_t_* are related. In other words, there is an edge connecting microbe j and microbe t in the microbial interaction network.

We have assumed that the relationship among microbes can be characterized by the multiclass Ising model (2)-(4). The state variables *Z_j_*’s in Ising model, however, are latent and can not be observed directly. Instead, the observable quantities are the relative abundances of the microbes which ae denoted by *X_j_*’s here. For each *X_j_*, we assume its distribution can be characterized by a mixture distribution which relies on the realization of **Z**. Specifically, we have the following conditional distribution for *X_j_* given *z_j_* = l for 1 ≤ *l* ≤ *K* – 1,

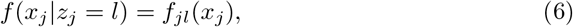

where *f_jl_* (1 ≤ l ≤ *K* – 1) can be any continuous distribution defined on [0,1]. While for *l* = 0 we have

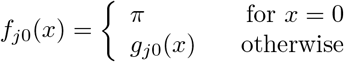

for some 0 − π − 1. Here *g*_*j*0_ can be any continuous distribution defined on [0,1]. In other words, *f*_*j*0_(*x*) is a zero-inflated distribution. Let *μ_jl_* = *E*(*X_j_* |*Z_j_* = l). For *l* = 0, *μ_jl_* is understood as the expectation with respect to the density function *g*_*j*0_. In order to ensure the model identifiability, we need the following assumption,

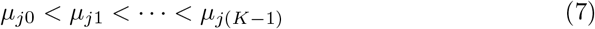

for 1 ≤ *j* ≤ *p*. Given **X** = (*X*_1_, …, *X_p_*) and its n i.i.d observations, **X**_1_, …, **X**_*n*_, we aim to estimate the MIN through (1)~(7) which we call zero-inflated latent Ising model (ZILI). The data generation process of ZILI is depicted in Figure 1.

**Figure 1:**
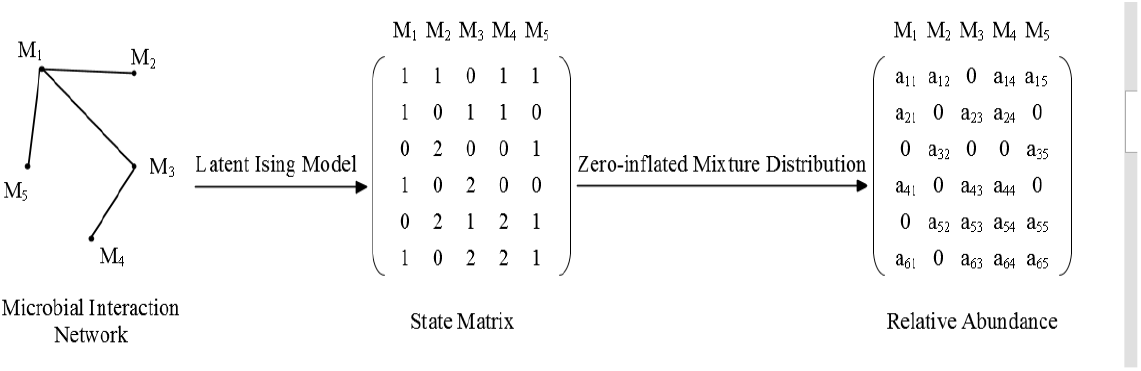
Diagram of data generation process for relative abundance of microbiota in ZILI model.

### Remark

We have adopted a zero-inflated form for density function *f*_*j*0_ while continuous form for *f_jl_* (1 ≤ l ≤ *K* – 1). In other words, the zero observations can only arise from *f*_*j*0_ which has the smallest mean relative abundance among *f*_*j*0_, …, *f*_*j*0(*K*–1)_. This assumption serves to ensure the identifiability of ZILI model. Note in literature, the zero observations in microbiome data are usually classified into two categories by their nature [7, 24]. In first category, the zero means the corresponding microbe physically does not exist in the subject, or true zero. In second category, the microbe does exist in the subject. Nevertheless, for this sample, this microbe happens not to exist or below the threshold of the testing procedure, *i.e.*, false zero. So our assumption about *f_jl_* for 0 ≤ l ≤ *K* – 1 means that both true and false zero’s can only come from *f*_*j*0_. Though there is possibility that this assumption may not hold in practice, we believe it is a reasonable approximation to the real situation.

## 3. Selection of MIN Based on ZILI Model

From equation (5), it can be seen that the selection of MIN is equivalent to the selection of the nonzero components of ***θ*** involved in ZILI model. In this section, we propose a two-step algorithm to select such nonzero components of ***θ*** based on **X**_1_, …, **X**_*n*_, the observations of relative abundance.

### 3.1. Step 1: state estimation

In this step, for each microbe, we aim to estimate the state *Z_j_* (1 ≤ *j* ≤ *p*) for each observation. Given the microbe, the proposed algorithm only involves its own observations. So for ease of exposition, we suppress the subscript *j* and use the generic notation (*Z, X*) to introduce the algorithm. The corresponding number of classes is denoted by *K_j_* = *K*.

With the observations of relative abundance, *X*_1_, *X*_2_, …, *X_n_*, in hand, the estimation of *Z* is carried out through the following optimal classification of *X*_1_, *X*_2_, …, *X_n_*. Without loss of generality, we assume that the observations have been ordered, i.e., *X*_1_ ≤ *X*_2_ ≤ … ≤ *X_n_*. For a given integer *k* ≥ 2, let *b*(*n, K*) denote a classification scheme which classifies (*X*_1_, …, *X_n_*) into *k* classes. Such classification can be depicted by the following notations,

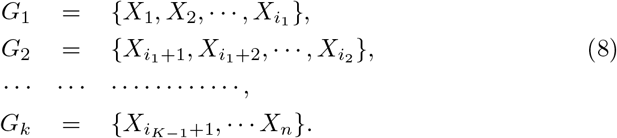

With notation *i*_0_ = 1, *i_k_* = *n*, we define the following loss function for *b*(*n, K*),

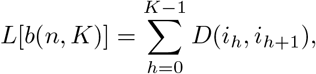

where

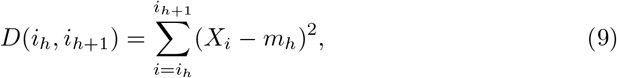

where

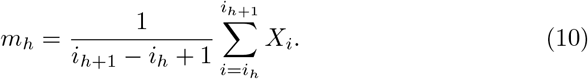

We aim to find a classification scheme *b*(*n, K*) which can minimize loss function *L*[*b*(*n, K*)]. Such optimal classification scheme is denoted by *p*(*n, K*). We employ the following top-down dynamic programming algorithm to find *p*(*n, K*)[40]. Specifically, the algorithm involves the following recursive procedures,

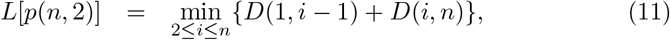

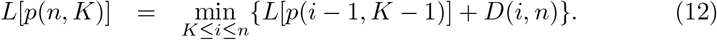

Based on (11)-(12), for given *K*, the algorithm can be implemented as follows. First, find *i*_*K*–1_ such that

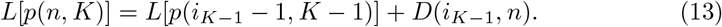

Based on *i*_*K*–1_, denote the *K* th class by *G_K_* = {*i*_*K*–1_, *i*_*K*–1_ + 1, …, *n*}. In second step, find *i*_*K*–2_ such that

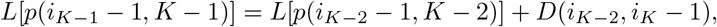

then we get the (*K* – 1)th class *G*_*K*–1_ = {*i*_*K*–2_, *i*_*K*–2_ + 1, …, *i*_*K*–1_ – 1}. By the same fashion, all the classes *G*_1_, *G*_2_, …, *G_K_* can be derived, which is the optimal solution *p*(*n, K*). Based on *p*(*n, K*), the estimate of *Z* for observations in class *G_k_* is defined as 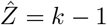 for *k* = 1, …, *K*.

The algorithm above assumes that *K*, the number of the classes, is known as a prior. In practice, *K* is usually unknown and has to be determined based on the data. Though several methods have been proposed in literature, such as likelihood ratio test in R package *mixtools* [43], or BIC method in package *sBIC* [48], these methods have poor performance when the data are zero-inflated. Instead, we propose the following criterion to select *K*. For a given upper bound, say, 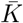, for each *K* with 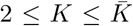, the minimum loss *L*(*p*(*n, K*)) is calculated. Define *d_K_* = *L*(*p*(*n, K* +1)) – *L*(*p*(*n, K*)) for 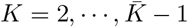 and let 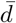 be the mean of *d_K_*’s. Then the first *K* with 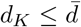 will be selected as the class number. This criterion turns out to have a better performance than the methods mentioned above in the simulation studies in Section 4.

### 3.2. Step 2: network selection

Equation (5) shows that, after the logit transformation, the conditional probability *P*{*Z_j_* = *l*|***Z***_*i*(−*j*)_ = **z**_*i*(−*j*)_} is a linear function of ***θ***. Here the covariates are the indicator functions of events { *Z_it_* = *h*} (*t* ≠ *j, h* = 1, …, *K* – 1). Based on this observation, the neighborhood method is proposed in [36] to select the nonzero components in ***θ*** for dichotomous Ising model. Here since *Z_ij_* is latent variable in ZILI, we replace *Z_ij_* by its estimate 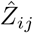 and then adopt the neighborhood method to select the MIN. Specifically, for *j*th microbe, let 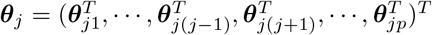 where ***θ***_*jt*_ is defined in Section 2. Based on the equation (5), we consider the following penalized group logistic regression problem,

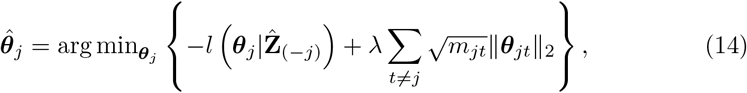

where 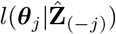 is the multinomial likelihood function, λ is the tuning parameter, *m_jt_* is the length of vector ***θ***_*jt*_ and || · ||_2_ is the Euclidean norm. The form of 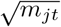 aims to account for the varying group size of ***θ***_*jt*_ [32]. Such form of penalty in (14) tends to shrink the components in same group **θ**_*jt*_ to zero simultaneously. For given λ, the coordinate decent algorithm [18, 19] is employed here to solve (14). As for the selection of λ, extended BIC proposed in [10] is adopted which favors sparser model compared with the standard BIC. The minimization problem in (14) is solved for each *Z_j_* (1 ≤ *j* ≤ *p*). With the final estimate 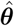 in hand, we define an edge between *Z_j_* and *Z_t_* if there exists at least one nonzero component in either 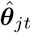 or 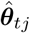. An alternative way to define an edge requires there exists at least one nonzero component in both 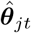 and 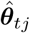. It turns out these two strategies are asymptotically equivalent [31, 36] and so we just employ the former one to select the MIN in the numerical studies. The magnitude of the components of 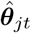 plays no role in the determination of the edges [11, 36].

In the above, the proposed algorithm estimates the interaction network by separately solving *p* conditional penalized maximum likelihood estimation problems. Alternatively, we can form a joint conditional likelihood function for ***θ*** and estimate the network by minimizing the penalized version of the joint conditional likelihood function. This approach, however, is not computationally as stable as (14) [11]. We therefore put the focus on the individual regression method (14). Figure 2 shows the workflow of the two-step algorithm through a simple artificial MIN.

**Figure 2:**
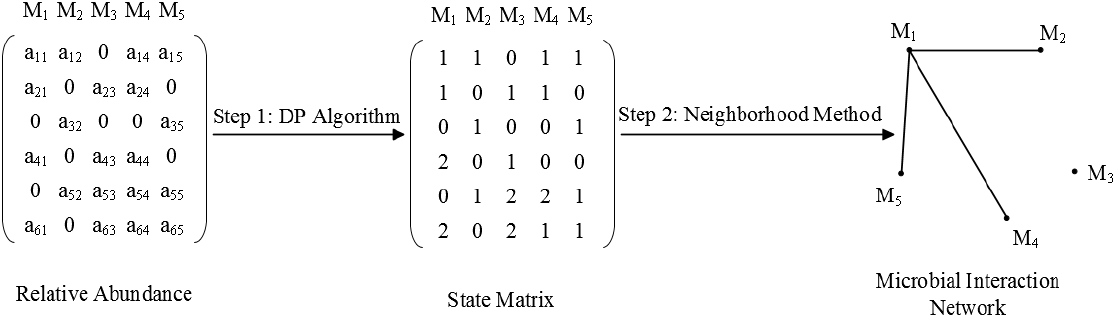
Diagram of two-step algorithm for network selection based on ZILI model.

#### Remark

For the two-step algorithm proposed above, it is expected that the selection of MIN will get improved if we can improve the state estimates 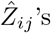. However, the misclassification is inevitable in two-step algorithm which will adversely affect the final network selection. In Section 4, we investigate how the misclassification impacts the MIN selection through simulation studies.

## 4. Simulation Studies

In this section, we investigate the performance of the two-step algorithm when ZILI is the underlying data generation model. As a comparison, the popular Gaussian graphical model (GGM) and dichotomous Ising model (DIS) will also be fitted using the same dataset. Here DIS is constructed by transforming the relative abundance into 0 or 1 according to whether it is less than the the median. The same algorithm in Section 3.2 will be employed to estimate the structure of this dichotomous Ising model.

Specifically, assume that there are *p* microbes with state variables **Z** = (*Z*_1_, …, *Z_p_*). Each realization of *Z_j_* (*j* = 1, …, *p*) takes value from the set {0,1,2}. The conditional distribution of *Z_j_* (*j* ≠ 1) given all the other components of **Z** only depends on microbe *Z*_*j*–1_. As for microbe 1, the distribution of *Z*_1_ depends on microbe *Z_p_*. For such a model, the nonzero parameters involved in equation (5) include (*θ*_*j*;1_, *θ*_*j*;2_, *θ*_*j*(*j*–1);11_, *θ*_*j*(*j*–1);12_, *θ*_*j*(*j*–1);21_, *θ*_*j*(*j*–1);22_) which are same for all *j*’s. For each repetition, these parameters are sampled from the multivariate normal distribution *N*_6_(*μ, Σ*) with *μ* = (– 1, 3, –0.8, 2, –3, –4)^*T*^ and Σ= diag(0.1^2^, 0.3^2^, 0.08^2^, 0.2^2^, 0.3^2^, 0.4^2^).

Given the Ising model above, the Gibbs sampler is employed to generate the samples of **Z**. Specifically, first a p-dimensional vector is generated where the states for each *Z_j_* are independently sampled from the set {0, 1, 2} with equal probability 1/3. Then given all *Z_t_*, (*t* ≠ *j*), the state of *Z_j_* is updated based on equation (5). By the same fashion, the states of all the other *Z_j_* can be updated recursively. We run this process 200 times and the final state of **Z** will be deemed a qualified representative of the underlying Ising model. Based on the samples of **Z**, the samples of absolute abundance **X** = (*X*_1_, …, *X_p_*) are generated according to *X_j_*|*Z_j_* = *z* ~ *N*(*μ_z_*,*σ*^2^) with *μ*_0_ = 10, *μ*_1_ = 15, *μ*_2_ = 20 and a given *σ*^2^. Pooling all the samples of **X** together leaves us a *n* × *p* matrix which represents n absolute abundance observations for *p* microbes. For each column, the absolute abundances which are less than a given percentile with rank *u* are replaced by zero. Here *u* is sampled from uniform distribution *U* [0, *z*] for a given 0 < *z* < 1. For each row in this zero-inflated matrix, we then transform the absolute abundances to relative abundances by dividing each entry by the corresponding sum of the row.

To compare the performances of different models, two criteria, true positive rate (TPR) and false positive rate (FPR) will be used which are defined respectively as,

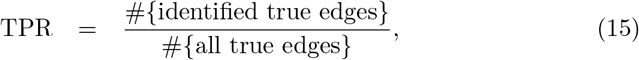

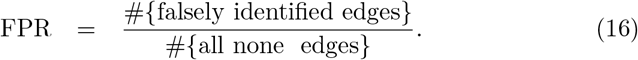

An ideal algorithm should have a relatively high TPR and low FPR. There are multiple factors that can influence the performance of the algorithm, which include the variance *σ*^2^, the sample size n, and the zero proportion *z*. For three choices *σ*, two choices of *n* and three choices of *z*, Table 1 lists the results of TPR and FPR for ZILI, DIS and GGM respectively. Here the number of the microbes is set to be *p* = 60 and the number of repetition is 100. Note for GGM, there are different estimation methods available such as graphical lasso [16], or neighborhood method [31] *et al.* Here in order to facilitate the comparison with ZILI and DIS, we adopt the neighborhood method of [31]. The same model selection criterion extended BIC is used in all cases. It can be seen from Table 1 that for all the scenarios considered, the proposed two-step algorithm does can select the network structure effectively while both GGM and DIS have very low TPR and can not properly select the true edges. On the other hand, all the three factors considered, i.e., variance, sample size and zero proportion have significant impact on the performances of two-step algorithm. Two-step algorithm has the best performance with the small *σ*^2^, *z* and large *n* which is in accordance with our expectation. In particular, a large *σ*^2^ will lead to a high misclassification rate for the state estimation in two-step algorithm which in turn results in a poor network selection, i.e., low TPR and high FPR.

**Table 1:** Comparison of ZILI, DIS and GGM for simulated data

|  |  |  | ZILI |  | DIS |  | GGM |  |
| --- | --- | --- | --- | --- | --- | --- | --- | --- |
| z | $\sigma$ | n | TPR | FPR | TPR | FPR | TPR | FPR |
| 10 | 0.5 | 60 | 0.8322 | 0.0110 | 0.0115 | 0.0008 | 0.1940 | 0.0347 |
|  |  | 120 | 0.9650 | 0.0024 | 0.0173 | 0.0006 | 0.0721 | 0.0058 |
|  | 1 | 60 | 0.7945 | 0.0123 | 0.0106 | 0.0007 | 0.0145 | 0.0059 |
|  |  | 120 | 0.9615 | 0.0043 | 0.0163 | 0.0005 | 0.1683 | 0.0051 |
|  | 2 | 60 | 0.2688 | 0.0256 | 0.0115 | 0.0010 | 0.0123 | 0.0061 |
|  |  | 120 | 0.5041 | 0.0187 | 0.0096 | 0.0003 | 0.0368 | 0.0046 |
| 40 | 0.5 | 60 | 0.7775 | 0.0134 | 0.0106 | 0.0009 | 0.1961 | 0.0274 |
|  |  | 120 | 0.9445 | 0.0059 | 0.0180 | 0.0005 | 0.1923 | 0.0056 |
|  | 1 | 60 | 0.7260 | 0.0139 | 0.0163 | 0.0007 | 0.1895 | 0.0276 |
|  |  | 120 | 0.9353 | 0.0067 | 0.0120 | 0.0003 | 0.1821 | 0.0057 |
|  | 2 | 60 | 0.2085 | 0.0247 | 0.016 | 0.0013 | 0.1328 | 0.030 |
|  |  | 120 | 0.4043 | 0.0194 | 0.0115 | 0.0004 | 0.0991 | 0.0055 |
| 80 | 0.5 | 60 | 0.4443 | 0.0217 | 0.0720 | 0.0086 | 0.2346 | 0.0465 |
|  |  | 120 | 0.6090 | 0.0148 | 0.1115 | 0.0071 | 0.2248 | 0.0144 |
|  | 1 | 60 | 0.4088 | 0.0214 | 0.0681 | 0.0085 | 0.2321 | 0.0477 |
|  |  | 120 | 0.6020 | 0.0149 | 0.1138 | 0.0071 | 0.2196 | 0.0146 |
|  | 2 | 60 | 0.1538 | 0.0247 | 0.0493 | 0.0084 | 0.1851 | 0.0514 |
|  |  | 120 | 0.2820 | 0.0207 | 0.0938 | 0.0065 | 0.1620 | 0.0142 |

## 5. Application to Infant Gut Microbiota

In this section, ZILI is employed to investigate the association among the microbes of microbiota in the infant stool sample from New Hampshire Birth Cohort Study (NHBCS), a cohort of mother-infant pairs in New Hampshire. For this dataset, stool samples were collected from infants at six weeks and twelve months of age, who were followed in the NHBCS. The stool samples were characterized by 16S rRNA sequencing. The R software package *DADA21* was used to infer the abundance of amplicon sequence variants in each sequenced sample [6]. Taxonomy at the family level was obtained by classifying the sequences against the reference training dataset from the GreenGenes Database Consortium (Version 13.8). There were 398 six week and 316 twelve months samples with varying abundances across 134 taxonomic families.

For each taxon, if the proportion of nonzero observations is less than 1%, then the number of classes is set to be *K_j_* = 2 and the observations are classified according to whether it is zero or not. Otherwise, the upper bound of *K_j_* is set to be 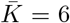. Then we follow the two-step algorithm to select the network. In order to gain insight from the difference between ZILI and GGM, the networks based on GGM have also been selected using the neighborhood method. In light of the severe zero inflation in the dataset, it is inappropriate to assume the GGM for the whole dataset. To alleviate the problem of zero inflation, we choose to use the subsets of this dataset to construct the GGM networks. Specifically, for each s = 10%, 20%, …, 80%, we extract the corresponding subset from the original dataset which only includes the microbes whose proportions of nonzero observations are greater than *s*. For each of these subsets, GGM is fitted using the neighborhood method. The ZILI network involves 134 microbe taxa while the eight GGM networks only involves eight subsets of these 134 taxa. So in order to compare the ZILI network with the eight GGM networks, we extract the subnetworks from ZILI networks for each *s*. For each of the extracted network, we then compare it with the corresponding GGM network in terms of their connectivity and the results are listed in Table 2.

**Table 2:** Comparison of microbial interaction networks selected by GGM and ZILI. The data are the relative abundances of microbiota in infant gut from NHBCS

|  |  | ZILI |  |  |  |  |
| --- | --- | --- | --- | --- | --- | --- |
|  |  | (0,0) | (1,0) | (0,1) | (1,1) | p-value |
| GGM | 10% | 589 | 58 | 6 | 13 | 0.0000 |
|  | 20% | 357 | 37 | 3 | 9 | 0.0000 |
|  | 30% | 182 | 38 | 2 | 9 | 0.0000 |
|  | 40% | 137 | 25 | 2 | 7 | 0.0000 |
|  | 50% | 89 | 23 | 2 | 6 | 0.0022 |
|  | 60% | 55 | 17 | 0 | 6 | 0.0005 |
|  | 70% | 46 | 15 | 0 | 5 | 0.0252 |
|  | 80% | 19 | 6 | 0 | 3 | 0.0445 |

|  |  | (0,0) | (1,0) | (0,1) | (1,1) | p-value |
| --- | --- | --- | --- | --- | --- | --- |
| GGM | 1% | 916 | 953 | 19 | 65 | 0.0000 |
|  | 2% | 1148 | 335 | 25 | 32 | 0.0000 |
|  | 3% | 816 | 482 | 31 | 49 | 0.0000 |
|  | 4% | 628 | 315 | 37 | 47 | 0.0000 |
|  | 5% | 503 | 403 | 32 | 52 | 0.0027 |
|  | 6% | 446 | 385 | 30 | 43 | 0.0198 |
|  | 7% | 270 | 466 | 21 | 63 | 0.0102 |
|  | 8% | 339 | 345 | 19 | 38 | 0.0192 |

In Table 2, each row corresponds to a pair of ZILI and GGM networks. For two microbes, (0,0) represents there is no edge connecting them in both ZILI and GGM network; (0,1) represents there is an edge in ZILI network while no edge in GGM network; (1,0) represents there is an edge in GGM network while no edge in Ising network; (1,1) represents there is an edge connecting them in both ZILI and GGM network. The columns 3-6 in Table 2 list the numbers of the edges falling into these four categories respectively. The relationship of ZILI and GGM is our primary interest. To this end, the *χ*^2^ test for the independence of ZILI and GGM is carried out and the corresponding adjusted p-value’s are listed in the last column of Table 2. Note the p-value here is based on the estimated networks rather than the relative abundance. So we call them conditional p-value. These p-value’s suggest that the networks of ZILI and GGM are closely related, even though ZILI and GGM are based on entirely different assumptions about how the data are generated. A more detailed inspection reveals that most of the edges selected by ZILI are also selected by GGM and GGM selects far more edges than ZILI. In other words, ZILI is more conservative than GGM in terms of edge selection.

Figure 3 presents the ZILI network and Figure 4 presents the GGM network corresponding to the threshold s = 10%. The other networks for *s* = 20%, …, 80% are available in the supplementary materials. Figure 5 presents the subnetwork that is shared by the networks in Figures 3 and 4. From Figure 5, it can be seen that Lachnospiraceae is selected as hub taxon by both ZILI and GGM. It has been discovered in literature that R. gnavus, one of the members in Lachnospiraceae family, has high frequency in infant gut [38]. Lachnospiraceae has close connections with severe human diseases, such as inflammatory bowel diseases (IBD) [35], non-alcoholic fatty liver disease [39]. The R. gnavus ATCC 29149 strain possesses the complete Nan cluster involved in sialic acid metabolism for the production of an intramolecular trans-sialidase [41]. It has also been demonstrated recently that R. gnavus produces iso-bile acids. The iso-bile acids detoxification pathway influences the growth of one of the predominant genera in the human gut, i.e., the Bacteroides [14]. In summary, Lachnospiraceae plays an active role in human metabolism which in turn may impact the growth of the other taxa in the gut microbiota. In this respect, it is not surprising to find its wide connections with other members of the microbiota.

**Figure 3:**
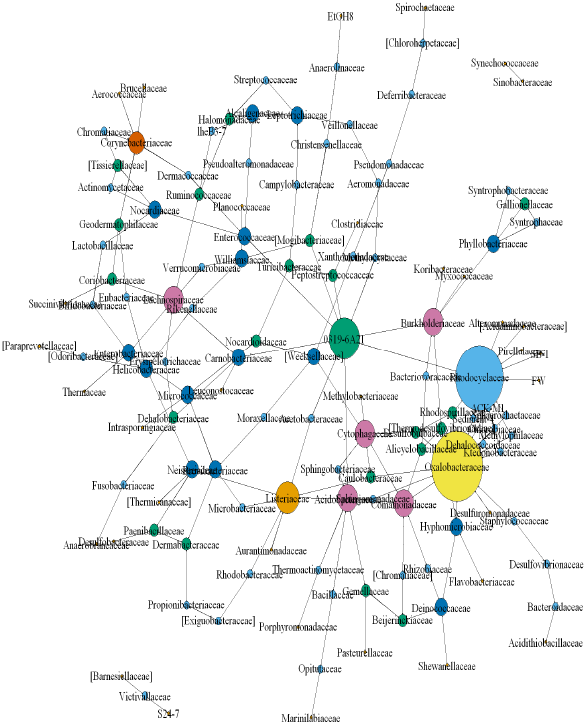
The network selected by ZILI for the microbiota in infant gut.

**Figure 4:**
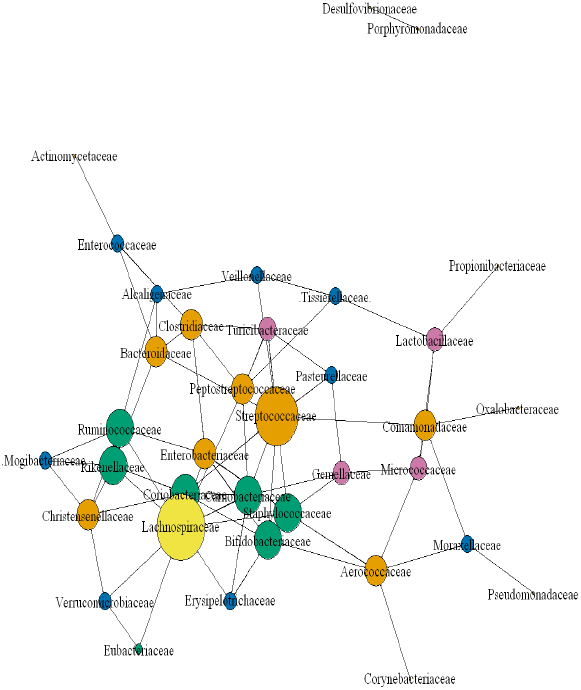
The network selected by GGM with threshold *s* = 10% for the microbiota in infant gut.

**Figure 5:**
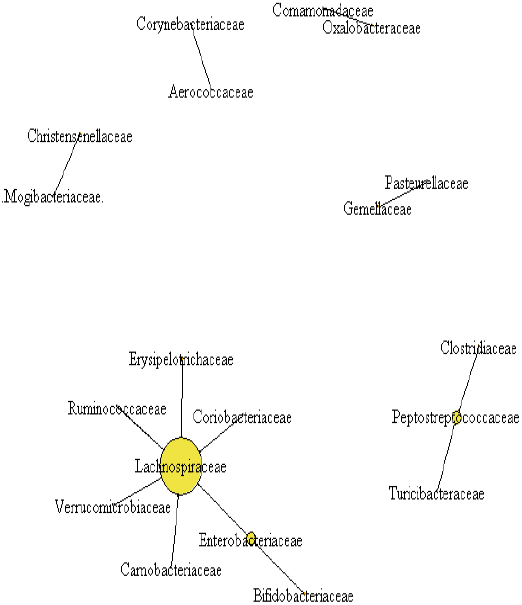
The subnetwork shared by the networks in Figures 3 and 4.

## 6. Discussion

The prosperous microbiome datasets have led us to a new level of biological researches. Nevertheless, how to gain scientific insight from these complex datasets through novel statistical methods remains a big challenge for researchers. In light of the difficulties in MIN selection, we propose a novel zero-inflated latent Ising model (ZILI) to this problem. In ZILI, the relative abundances of microbiota are assumed to follow a mixture distribution which relies on the realization of a latent Ising model. Through simulation studies, it is shown that under given scenarios, the proposed two-step algorithm for the inference of ZILI can select the true network structure effectively while Gaussian graphical model and dichotomous Ising model have little power to recover the network structure. For a microbiome dataset from New Hampshire Birth Cohort Study, it is shown that ZILI is more conservative compared with Gaussian graphical model. Among the edges shared by these networks, a hub taxon is selected which has close connections with human metabolism. These findings indicate that ZILI can serve as an competitive model to estimate the microbial interaction network. On the other hand, given the statistically significant overlap between ZILI and GGM networks, it is interesting to investigate the performance of ZILI and its relationship with traditional methods for the microbiota in other body parts in the future studies.

## Acknowledgements

We are grateful to the participants and staff of the New Hampshire Birth Cohort Study for providing the processed microbiome data. This work is supported in part by US National Institutes of Health grants (R01LM012012; R01LM012723; P20ES018175; P01ES022832; UG3OD023275), US Environmental Protection Agency grant (RD83459901), the Children’s Center grant, the Superfund grant and COBRE.

## Notes

### Competing Interest Statement

The authors have declared no competing interest.

